# Ultrastructure expansion microscopy of *Plasmodium* gametocytes reveals the molecular architecture of a microtubule organisation centre coordinating mitosis with axoneme assembly

**DOI:** 10.1101/2021.07.21.453039

**Authors:** Ravish Rashpa, Mathieu Brochet

## Abstract

The transmission of malaria-causing parasites to mosquitoes relies on the production of the gametocyte stages and their development into gametes upon a blood feed. These stages display various microtubule cytoskeletons and the architecture of the corresponding microtubule organisation centres (MTOC) remains elusive. Combining ultrastructure expansion microscopy (U-ExM) with bulk proteome labelling, we first reconstructed in 3D the subpellicular microtubule network and its associated actin cytoskeleton, which confer cell rigidity to *Plasmodium falciparum* gametocytes. Upon activation, as the microgametocyte undergoes three rounds of endomitosis, it simultaneously assembles axonemes to form eight flagellated microgametes. Here, U-ExM combined with Pan-ExM revealed the molecular architecture of a single bipartite MTOC coordinating mitosis with axoneme formation. This MTOC spans the nuclear membrane linking acentriolar mitotic plaques to cytoplasmic basal bodies by proteinaceous filaments. The eight basal bodies are concomitantly *de novo* assembled from a deuterosome-like structure, where centrin, γ-tubulin, SAS4/CPAP and SAS6 form distinct subdomains. Once assembled, the basal bodies show a fusion of the proximal and central cores where colocalised centrin and SAS6 are surrounded by a SAS4/CPAP-toroid in the lumen of the microtubule wall. Sequential nucleation of axonemes and mitotic spindles is associated with a dynamic movement of γ-tubulin from the basal bodies to the acentriolar plaques. We finally show that this atypical MTOC architecture relies on two non-canonical MTOC regulators, the calcium-dependent protein kinase 4 and the serine/arginine-protein kinase 1. Altogether, these results provide insights into the molecular organisation of a bipartite MTOC that may reflect a functional transition of a basal body to coordinate axoneme formation with mitosis.

## Introduction

Malaria is caused by intracellular parasites of the *Plasmodium* genus that are transmitted via the bite of an infected Anopheles mosquito. The major pathophysiological processes in malaria are linked to the proliferation of asexual blood stages, whereas transmission to the mosquito is solely initiated by an obligatory sexual life cycle phase. Differentiation from asexually replicating stages into non-dividing transmission stages, the gametocytes, takes place inside erythrocytes. Following a period of maturation, micro- and macrogametocytes are ready to initiate transmission when ingested by a mosquito.

The biology of gametocytes shows contrasting features across *Plasmodium* species. *Plasmodium* gametocytes of a majority of species are round and mature in a day or two, whereas gametocytes of *P. falciparum*, the species responsible for most malaria-related human deaths, are sickle shaped and reach full morphological maturity in 8 – 12 days [1]. *P. falciparum* gametocytes show a development that is divided into five stages based on morphological changes observed by light microscopy or ultrastructural analyses [2]. In particular, transition to stage II is marked by the formation of a cisternal membrane structure just beneath the plasma membrane called the inner membrane complex (IMC). The IMC acts as a scaffold supporting a dense network of microtubules. By stage IV, the gametocytes are maximally elongated and display pointed ends. Upon transition to stage V, the microtubule network is disassembled and the extremities become more rounded [2].

Terminally differentiated gametocytes resume their development in the mosquito midgut following a blood meal, activated by the presence of xanthurenic acid (XA), a rise in pH, and a drop in temperature [3]. Upon activation both micro- and macrogametocytes increase in size and round up [4]. Egress from the host erythrocyte relies on the release of secretory vesicles called osmiophilic bodies [5], which are abundant in the macrogametocyte [6]. Following emergence and activation of translationally repressed mRNAs [7], macrogametes are rapidly available for fertilisation [8]. In contrast, microgametocyte resuming development completes three rounds of genome replication and endomitosis within a single nucleus. It simultaneously assembles eight axonemes to form eight flagellated microgametes in a process called exflagellation [9, 10].

Observations by electron microscopy (EM) gave general insights into the dynamics of microtubule during microgametogenesis [4]. The mature microgametocyte shows two electron-dense structures that have been linked with the formation of the mitotic spindles and the axonemes, respectively. One of this structure was described as an amorphous MTOC lying on the cytoplasmic face of a nuclear pore that is physically linked to another electron dense aggregation called the intranuclear body in the nuclear face of the same pore [9]. Upon activation of gametogenesis, the amorphous MTOC gives rise to eight basal bodies on which axonemes are nucleated [4], while the intranuclear body likely corresponds to the so-called centriolar plaques (also called spindle poles or centrosomes). The basal bodies of the forming axonemes remain attached to their respective centriolar plaque during the three following mitoses and are moved around the nuclear envelope at each round of division. After 1-2 min, the first mitotic spindle is visible and four basal bodies with nucleating axonemes are found in tetrad at each of the two opposing centriolar plaques. Three to five minutes later, following the assembly of the two spindles of mitosis II, two basal bodies lie at each end of the four centriolar plaques. By 6-8 min, the four spindles of mitosis III have formed and a single basal body is found at each of the eight centriolar plaques. Chromatin condensation only sets in at the end of mitosis III, when each of the eight short spindles anchors one haploid set of 14 chromosomes to each of the centriolar plaques. In *Plasmodium* gametocytes, axoneme polymerisation is intracytoplasmic and independent of intra-flagellar transport [11, 12]. Axonemes reach around 14 μm in length in just 10 min and lay coiled around the membrane of the nucleus. They display a classical organisation with nine doublets of microtubules arranged in a circular pattern surrounding a central pair of singlet microtubules [11]. At the onset of exflagellation, the axonemes become motile and swim basal body first out of the parental cell forming the flagellum [4]. The prior attachment of each basal body to a centriolar plaque allows each axoneme to drag a haploid genome into the forming flagellated gamete [13].

The exact biogenesis, structure, and composition of the basal body and the centriolar plaque in *Plasmodium* gametocytes remain unclear. Available data on the structural organisation and the biogenesis of the centriolar plaque is very limited. In asexual stages, the centriolar plaque shows a bipartite organization, with an extranuclear region containing centrin and an intranuclear DNA-free region harbouring microtubule nucleation sites [14]. It appears as an electron-dense area that neither shows a centriole nor any other known distinct structures such as the yeast spindle pole body, the *Dyctiostelium* nucleus associated body or the diatom microtubule centre. In the absence of centriole in the so-called centriolar plaque, we will hereafter refer to the acentriolar plaque to avoid any confusion. In gametocytes, the acentriolar plaques are linked to the basal bodies through nuclear pores or fenestra. While little is known about the acentriolar plaque and its molecular link to the basal body, the general organisation of the basal body is slightly less elusive. It is around 0.25 µm long. The central pair of axonemal singlet microtubules does not extend into the basal body but is underlaid by an electron-dense region where γ-tubulin was suggested to reside [15]. Radial spokes have been described between this mass and the peripheral microtubules. Despite coding for four centrins, the *Plasmodium* genomes lack many conserved components of the basal body except SAS6, SAS4/CPAP and CEP135 [16–18]. A *P. berghei* SAS6-KO clone displayed reduced basal body numbers, axonemal assembly defects and abnormal nuclear allocation [19].

Our understanding of the ultrastructural organisation of *Plasmodium* gametocytes has initially relied on EM. It was more recently complemented by fluorescence light microscopy or super-resolution microscopy using multiple markers to infer the dynamic and the molecular composition of observed structures albeit at a lower resolution. We have recently implemented U-ExM to image the *Plasmodium* cytoskeleton leading to the identification of a remnant conoidal structure in ookinetes [20]. Here we used U-ExM and Pan-ExM to visualise ultrastructural changes during gametocyte development and gametogenesis in both *P. falciparum* and *P. berghei* [20]. This allowed us to observe multiple cellular structures that were previously only accessible by electron or super resolution microscopy. In particular, by localising known markers of the basal body, we could refine the description of the structural and molecular organisation for this organelle and its link with the acentriolar plaque. We further show the differential requirement of two kinases in the homeostasis of this atypical bipartite MTOC.

## Results

### U-ExM of developing *P. falciparum* gametocytes highlights the subpellicular microtubule network

*P. falciparum* gametocytes show five distinct morphological stages during their development. The fine ultrastructural architecture of each stage has been previously thoroughly investigated by detailed conventional EM [4] or more recently by Serial Block Face Scanning EM [21]. Here we combined U-ExM with bulk proteome [22, 23] and α/β tubulin labelling to image the intracellular development of gametocytes using the iGP2 line [24].

The hallmark of stage II transition is the deposition of the IMC and a dense network of MT below the parasite plasma membrane. At this stage, NHS-ester labelling allowed discriminating the host erythrocyte showing a lighter density compared with the gametocyte. Inside the gametocyte, NHS-ester dense puncta corresponding to osmiophilic bodies were already observed in female gametocytes (Fig. 1A). Gametocytes showed a single pointed-end associated with an NHS-ester dense area. This region may correspond to a MTOC as suggested by α/β tubulin staining that shows a large ribbon of bundled microtubules below the parasite pellicle that loops across the cytoplasm. In stage III gametocytes, the usual lemon shaped organisation was observed, as well as the lateral expansion of the microtubule network. This expansion may coincide with the growth of the previously described 13 Phil1-positive IMC plates [21], however, the plates were not visible by NHS-ester staining at this stage (Fig. 1B).

**Figure 1.**
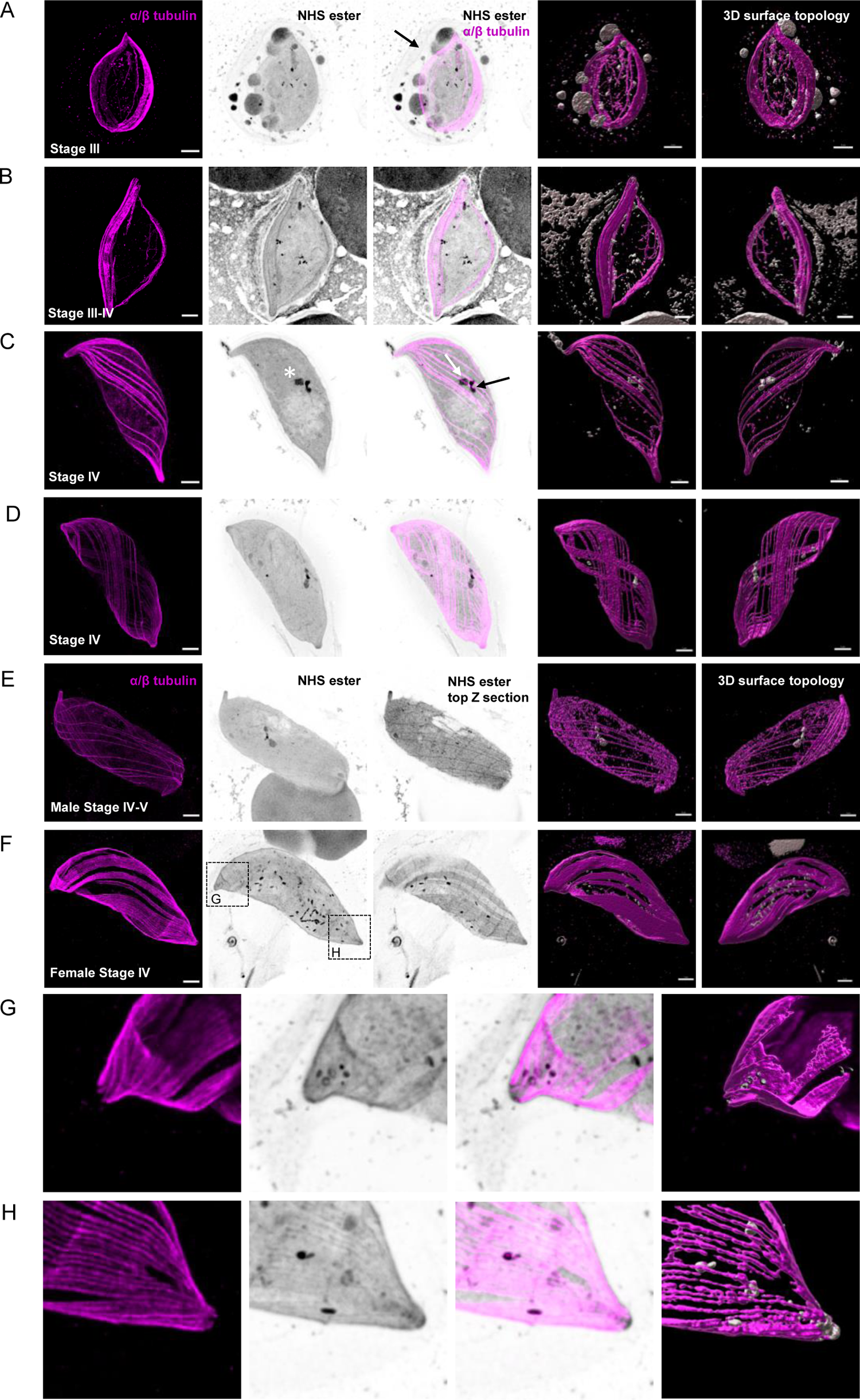
U-ExM allows reconstruction of *P. falciparum* gametocytogenesis in 3D. **A-F.** Representative full projections of *P. falciparum* gametocyte stages. α/β tubulin: magenta; amine reactive groups / NHS-ester: shades of grey. Columns 4 and 5 represent the 3D surface topology reconstruction of α/β tubulin and NHS-ester staining. **A.** A stage III lemon shaped gametocyte. The black arrow shows the host erythrocyte. **B.** A stage III to IV transitioning macrogametocyte, the tips start to get pointed and the cell displays a more elongated shape. Osmiophilic bodies show strong NHS-ester staining. **C and D.** A stage IV microgametocyte characterised by the presence of a cytoplasmic amorphous MTOC (black arrow) and the intra nuclear body (white arrow). The nuclear contour is distinguished by a light NHS-ester staining and the white asterisk indicates the position of the nucleus. Subpellicular microtubules start to surround the gametocyte. **E and F.** Stage IV to V micro- and macrogametocytes displaying rounded ends and an increase in width. Subpellicular microtubules start disassembling but IMC plates remain visible by NHS-ester staining (top Z section). In macrogametocytes, NHS-ester dense osmiophilic bodies are visible. At the gametocyte extremities (dotted area) 3 to 4 ring-like structures are present. **G and H**. Close-up images of the apical ends highlighting NHS-ester dense annuli. Scale bars = 5 µm.

By stage IV, the gametocytes are maximally elongated and display two pointed ends (Fig. 1C and D). Between these pointed ends, α/β tubulin staining showed continuous and evenly distributed microtubules running the length of the gametocyte with tightly packed arrays at both extremities. A high NHS-ester density additionally highlighted sutures between the 13 IMC plates suggesting a high protein density between plates (Fig. 1E and F). Conversely, a low NHS-ester density was observed adjacent to the nucleus. This may correspond to a membrane dense region such as the mitochondrion [2]. Similarly, the nuclear periphery was also highlighted by a lighter NHS-ester staining. The sexual dimorphism became most apparent at this stage: a NHS-ester dense structure spanning the nuclear membrane was observed in all microgametocytes. On the cytoplasmic side, it corresponds to the previously described amorphous electron dense MTOC from which axonemes nucleate (see below) (Fig. 1C-E). On the nuclear side this might correspond to the future acentriolar plaques that stain negative for DNA, as observed in asexual blood stages [14]. Upon transition to stage V, the microtubule network progressively disassembled and the tips sequentially became more rounded. As previously described in macrogametocytes, a large number of NHS-ester dense puncta corresponding to osmiophilic bodies were clearly visible (Fig. 1F). Intriguingly, NHS-ester dense annuli structures were also observed at both tips of female gametocytes (Fig. 1G and H).

### U-ExM of developing *P. falciparum* gametocytes highlights the complementary of the actin cytoskeleton with the subpellicular microtubules

NHS-ester staining highlighted denser fibres at the extremities of the subpellicular microtubules. We thus wondered whether these structures might correspond to the F-actin cytoskeleton beneath the IMC that was previously described [25]. An antibody against actin [26] strongly labelled the ends of stage IV micro-and microgametocytes where microtubules did not extend (Fig. 2). Labelling also extended lengthwise into the parasite body alongside the subpellicular microtubules as previously described [25].

**Figure 2.**
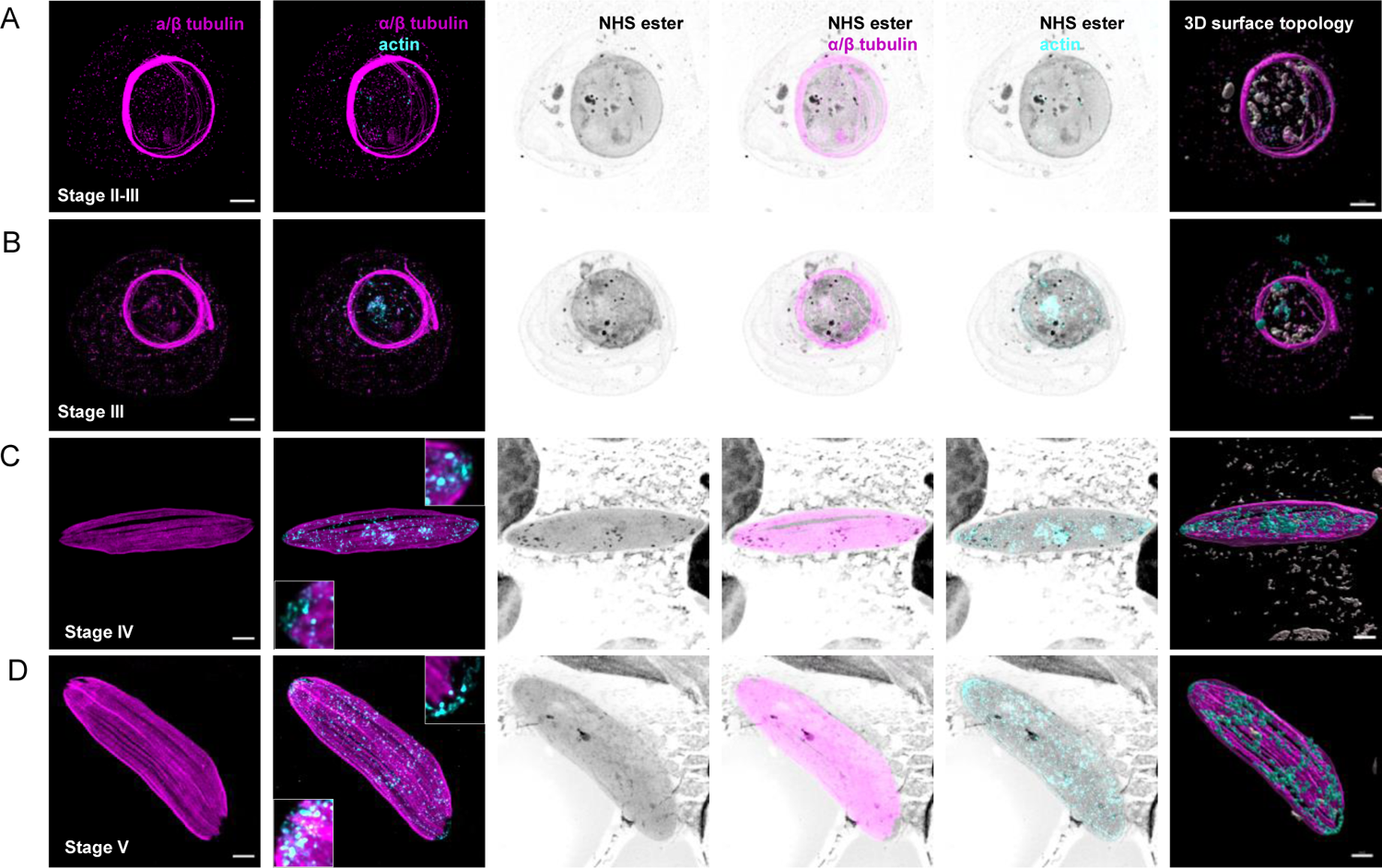
U-ExM confirms the organisation of the actin cytoskeleton during *P. falciparum* gametocytogenesis. **A-D**. Representative full projections of *P. falciparum* gametocyte stages. α/β tubulin: magenta; amine reactive groups / NHS-ester: shades of grey; actin: cyan. Column 6 shows the 3D surface topology reconstruction of α/β tubulin and actin. **A and B.** At early stages, a punctate distribution of actin is observed. **C-D.** From stage IV gametocytes, actin shows a polarised localisation, mostly concentrated at both extremities in tubulin-free areas showing a mesh-like organization. Actin also extends lengthwise alongside the subpellicular microtubules. Insets in (C and D) are close-up images of both extremities. Scale bars = 5 µm.

### 3D reconstruction of *P. falciparum* microgametogenesis

We then investigated the development of *P. falciparum* gametocytes into gametes upon activation with xanthurenic acid at a permissive temperature. Two to three minutes post-activation the remaining subpellicular microtubules were further degraded and the sutures between IMC plates were not distinguishable anymore (Fig. 3A and B). At this stage the surrounding erythrocyte was still visible. In macrogametocytes, osmiophilic bodies moved toward the periphery of the cell. No fusion of the vesicles with the parasite plasma membrane were seen by NHS-ester staining (Fig. 3A). In microgametocytes, the mitotic spindle was revealed by both NHS-ester and α/β tubulin labelling while the NHS-ester staining showed dots localising on the spindle that likely correspond to each of the 28 kinetochores (Fig. 3B). At both extremities of the spindle, two flat acentriolar plaques were strongly labelled by NHS-ester and no overlap with this structure was observed with the tubulin-positive spindle. On the cytoplasmic face of each acentriolar plaque, four NHS-ester dense basal bodies were associated with orthogonally arranged short axonemes (Movie S1). By ten minutes, microgametocytes have rounded up, undergone three rounds of endomitosis and assembled eight axonemes that are labelled by both NHS-ester and α/β tubulin (Fig. 3C).

**Figure 3.**
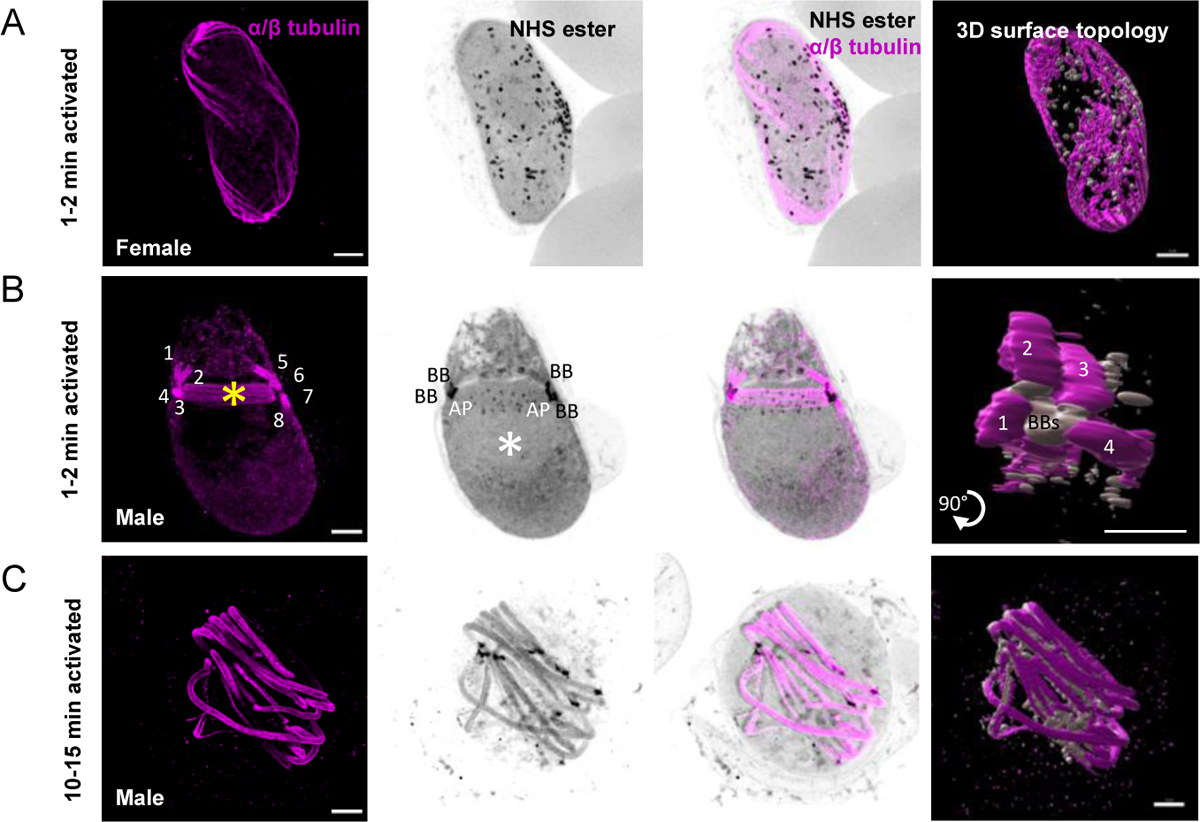
Mitotic structures and axonemes are highlighted by U-ExM during *P. falciparum* gametogenesis. **A-C.** Representative full projections of activated *P. falciparum* gametocytes. α/β tubulin: magenta; amine reactive groups / NHS-ester: shades of grey. Column 4 shows the 3D surface topology reconstruction of α/β tubulin and NHS-ester. **A.** A 1-2 min activated macrogametocyte shows degrading the subpellicular microtubule and NHS-ester dense osmiophilic bodies. **B.** A 1-2 min activated microgametocyte shows the NHS-ester dense basal bodies (BB) giving rise to axonemes (1 to 8 visible in supplementary movie 1). The mitotic spindle (yellow asterisk) is highlighted by α/β tubulin staining and, at each extremity, the acentriolar plaques (AP) show an NHS-ester dense staining. NHS-ester positive kinetochores are present along the spindle. At this stage, the subpellicular microtubules are completely lost. **C.** A 10-15 min activated microgametocyte displays a round shaped and full length axonemes. Scale bars = 5 µm.

### 3D reconstruction of *P. berghei* microgametogenesis highlights the spatial dynamic of the bipartite MTOC coordinating axoneme formation and mitosis

We then applied U-ExM to observe gametogenesis in *P. berghei* gametocytes. These have been widely used to investigate the molecular and cellular biology of *Plasmodium* transmission stages. Terminally differentiated *P. berghei* gametocytes were significantly different from their *P. falciparum* counterparts, displaying a round shape with no subpellicular microtubules nor IMC. While α/β tubulin labelling did not highlight any particular cellular structure at this stage, NHS-ester staining allowed visualising the host erythrocyte and delineating the contour of the nucleus with a lighter staining. To confirm that the nuclear membrane corresponded to a lighter NHS-ester, we took advantage of the previously published PbGEX1-HA line [27]. GEX1 was shown to be an ancient nuclear envelope protein family, essential for sexual reproduction in eukaryotes. GEX1-HA labelling colocalised with the lighter NHS-ester staining and highlighted a highly lobulated organisation of the nucleus in microgametocytes (Fig. 4A). NHS-ester staining also highlighted sexual dimorphism; macrogametocytes showed NHS-ester dense osmiophilic bodies (Fig. 4B).

**Figure 4.**
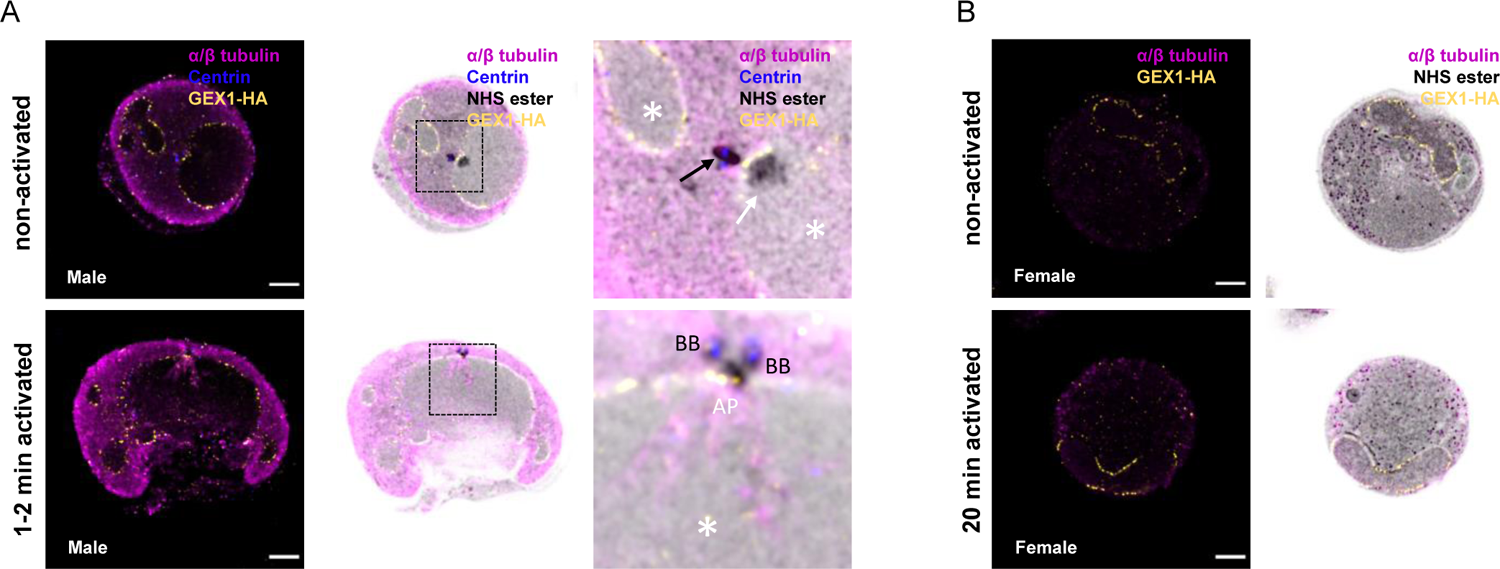
The MTOC coordinating axoneme formation and mitosis shows a bipartite structure across the nuclear membrane during *P. berghei* microgametogenesis. A and B. Localisation of the nuclear membrane protein GEX1-HA in *P. berghei* gametocytes. α/β tubulin: magenta; amine reactive groups/NHS-ester: shades of grey; centrin: blue; GEX1-HA: yellow. **A.** Non-activated and activated microgametocytes show a lobulated nuclear architecture, as seen by GEX1-HA signal and tubulin negative areas. Column 3 represents a close-up image of the boxed areas. The nuclear contour is distinguished by a light NHS-ester staining, which matches GEX1-HA staining. Nucleus: white asterisk; amorphous MTOC: black arrow (centrin-positive); basal bodies: BB (centrin-positive); Intranuclear body: white arrows (centrin-negative); acentriolar plaque: AP (centrin-negative). **B.** Non-activated and activated macrogametocytes. Macrogametocytes display a crescent-shaped nucleus, very low levels of α/β tubulin, and NHS-ester dense osmiophilic bodies. Scale bars = 5 µm.

*P. berghei* microgametocytes displayed a bipartite NHS-ester dense structure corresponding to the amorphous MTOC that was partially labelled by an antibody against human Centrin 1. This amorphous MTOC faced the NHS-ester positive intranuclear body that lies on the other side of the nuclear membrane (Fig. 5A). Upon activation by xanthurenic acid at a permissive temperature, the cell cycle events of the *P. berghei* gametocyte were comparable to *P. falciparum* with the sequential segregation and migration of 2×4, 4×2 and 8×1 pairs of acentriolar plaques and centrin-positive basal bodies from which eight axonemes assemble and grow (Fig. 5B to D). Upon exflagellation individual male gametes were observed with a strong NHS and centrin-positive basal body at one extremity and a diffuse Hoechst staining of DNA along the axoneme (Fig. 5E).

**Figure 5.**
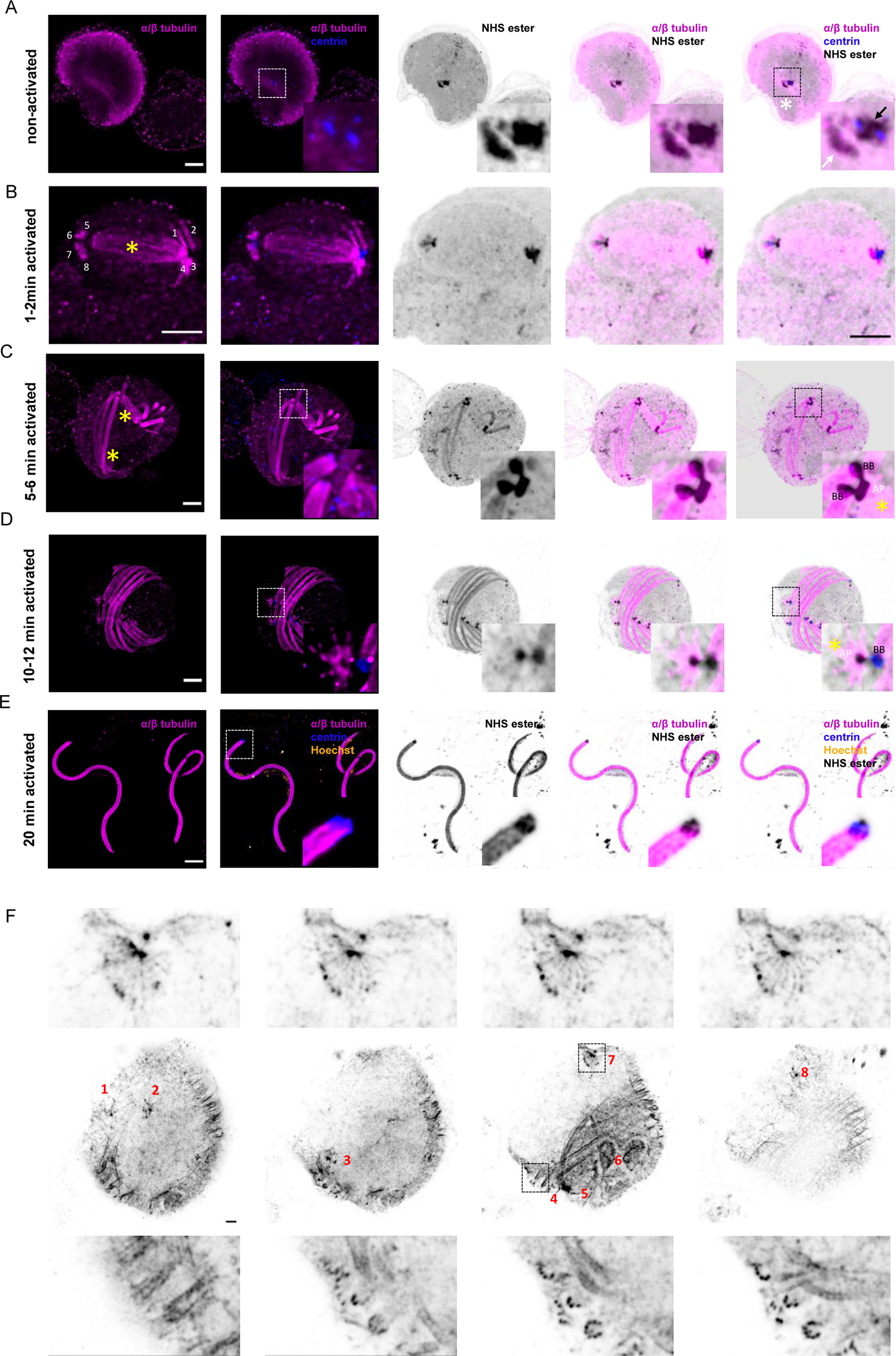
U-ExM and Pan-ExM of *P. berghei* microgametogenesis allow further insights into the dynamics of mitotic structures and axoneme formation. A-E. Representative full projections of activated *P. berghei* gametocytes. α/β tubulin: magenta; amine reactive groups/NHS-ester: shades of grey; centrin: blue. Boxed areas indicate areas shown in insets. **A.** A non-activated microgametocyte showing an amorphous MTOC (black arrow, centrin-positive) and intranuclear body (white arrow, centrin-negative). **B.** A 1-2 min activated microgametocyte, showing two tetrads of basal bodies and eight growing axonemes (1 to 8). In the nucleus the mitotic spindle is highlighted by a yellow asterisk. **C.** A 5-6 min activated microgametocyte is undergoing mitosis II with two mitotic spindles (yellow asterisks). Each acentriolar plaque (AP) is connected to two cytoplasmic basal bodies (BB). **D.** A 10-12 min activated microgametocyte has undergone mitosis III. Each acentriolar plaque (AP) is connected to a single cytoplasmic basal body (BB). The yellow asterisk highlights a remnant mitotic spindle. **E.** Exflagellated male gametes display a centrin (blue) and NHS-ester dense basal body at their proximal extremity while DNA (Hoechst: yellow) runs along the axoneme. **F.** Pan-ExM of a 10-12 min activated and NHS-ester labelled microgametocyte. The central panels represent four different z sections of the same microgametocyte. Each mitotic spindle is number from 1 to 8. Top panels show close-up images of four different z sections of mitotic spindle 7. Lower panels show four different z sections of the boxed area highlighting individual axonemal microtubules. Scale bars = 5 µm.

We then asked whether NHS-ester labelling of 2-times expanded cells (Pan-ExM), would provide further spatial resolution of the structures highlighted by U-ExM and NHS-ester staining, as previously shown for other Hela cells [23]. Sequential expansion allowed to expand *P. berghei* gametocytes 10.0 times. In 15 min activated gametocytes, Pan-ExM allowed resolving individual microtubules of the axonemes (Fig. 5F) and of the mitotic spindle where kinetochores are also NHS-ester positive. Interestingly, NHS-ester positive filaments linking the acentriolar plaque and its associated basal body were additionally observed. These proteinaceous filaments may correspond to the structure supporting the cohesion between the acentriolar plaque and the basal body during microgametogenesis.

### Molecular organisation of a bipartite MTOC made of basal bodies and acentriolar plaques

To get further insights into the ultrastructural organisation and relationship of the basal bodies and the acentriolar plaques, we set out to map possible markers of these structures on the NHS-ester dense labelling. We first focused on γ-tubulin, which is an indispensable component of MTOCs that contributes to microtubule nucleation and organisation in all eukaryotes (Fig. 6) [28]. It is recruited to the centrosome in metazoans or is tethered to the inner and outer plaques of the yeast spindle pole body. Additionally, we 3xHA-tagged two centriolar proteins (Fig. S1A-D): the proximal protein SAS6 that initiates the cartwheel assembly [29] and the proximal protein SAS4 that surrounds the centriole and facilitates the formation of the microtubule wall [30].

**Figure 6.**
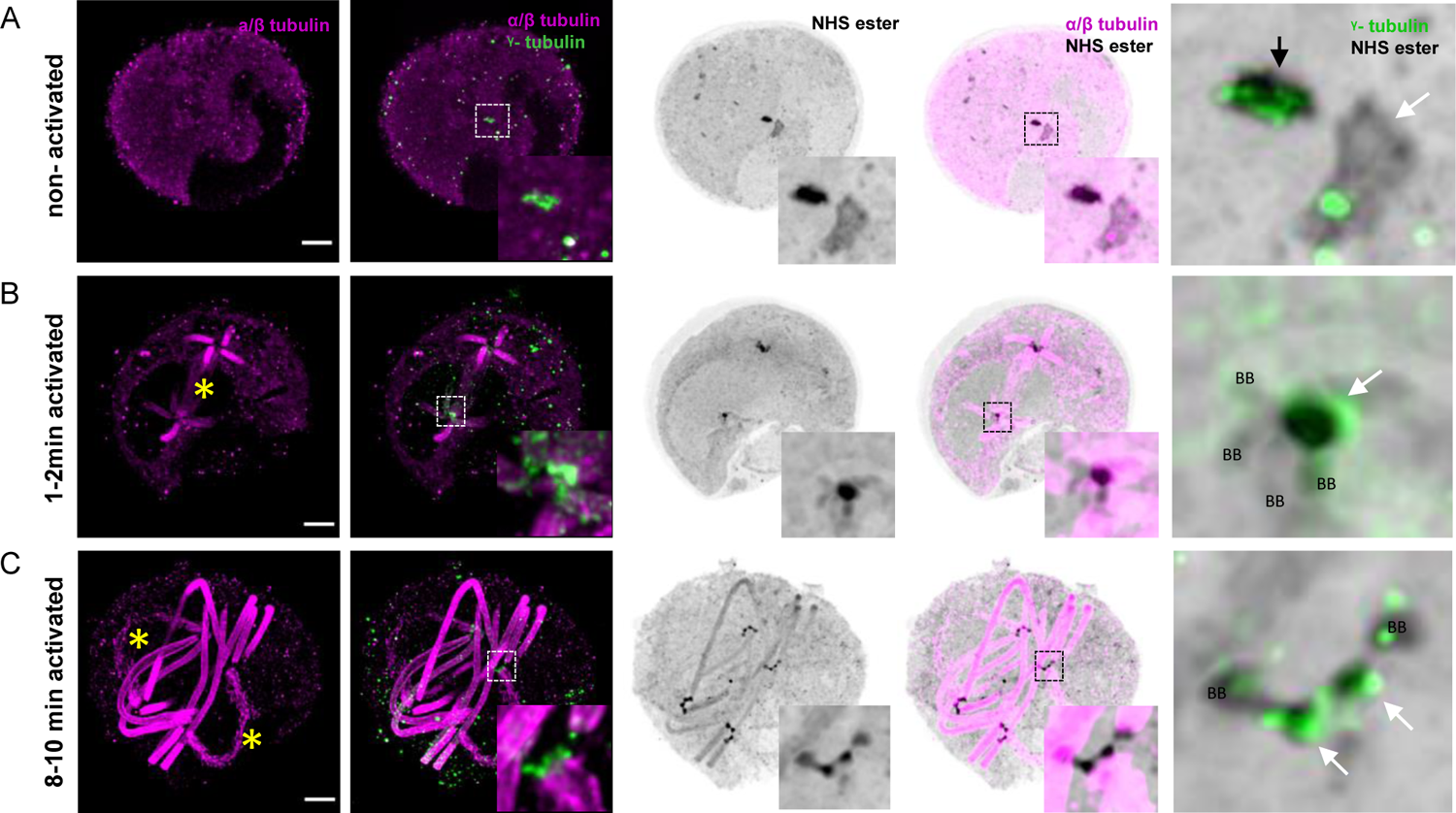
γ-tubulin displays dynamic localisation from the amorphous MTOC to basal bodies and acentriolar plaques during *P. berghei* microgametogenesis. A-C. Representative full projections of activated *P. berghei* gametocytes. α/β tubulin: magenta; amine reactive groups/NHS-ester: shades of grey; centrin: blue. Boxed areas indicate areas shown in insets. Column 5 represents zoomed-in images highlighting the basal bodies (BB), the amorphous MTOC (black arrow), the intranuclear body (white arrow) and acentriolar plaques (AP) stained with NHS-ester and for γ-tubulin. The yellow asterisks highlight mitotic spindles. Scale bars = 5 µm.

Prior to the activation of terminally differentiated microgametocytes γ-tubulin, SAS4-HA and SAS6-HA were only detected in the amorphous MTOC but not in the intranuclear body (Fig. 6A, Fig. 7A and E). At this stage we could not observe structures resembling basal bodies as previously described [4]. However, surface topology modelling indicated a certain level of organisation within the amorphous MTOC, where SAS4-HA was distributed at the periphery of the NHS-ester dense structure (Fig. 7A and Fig. 8A) in which SAS6-HA spheroids were distributed around centrin-positive regions (Fig. 7E and Fig. 8B). These observations indicate that despite the absence of basal bodies in non-activated gametocytes, their molecular components already display specific localisation patterns within the amorphous MTOC.

**Figure 7.**
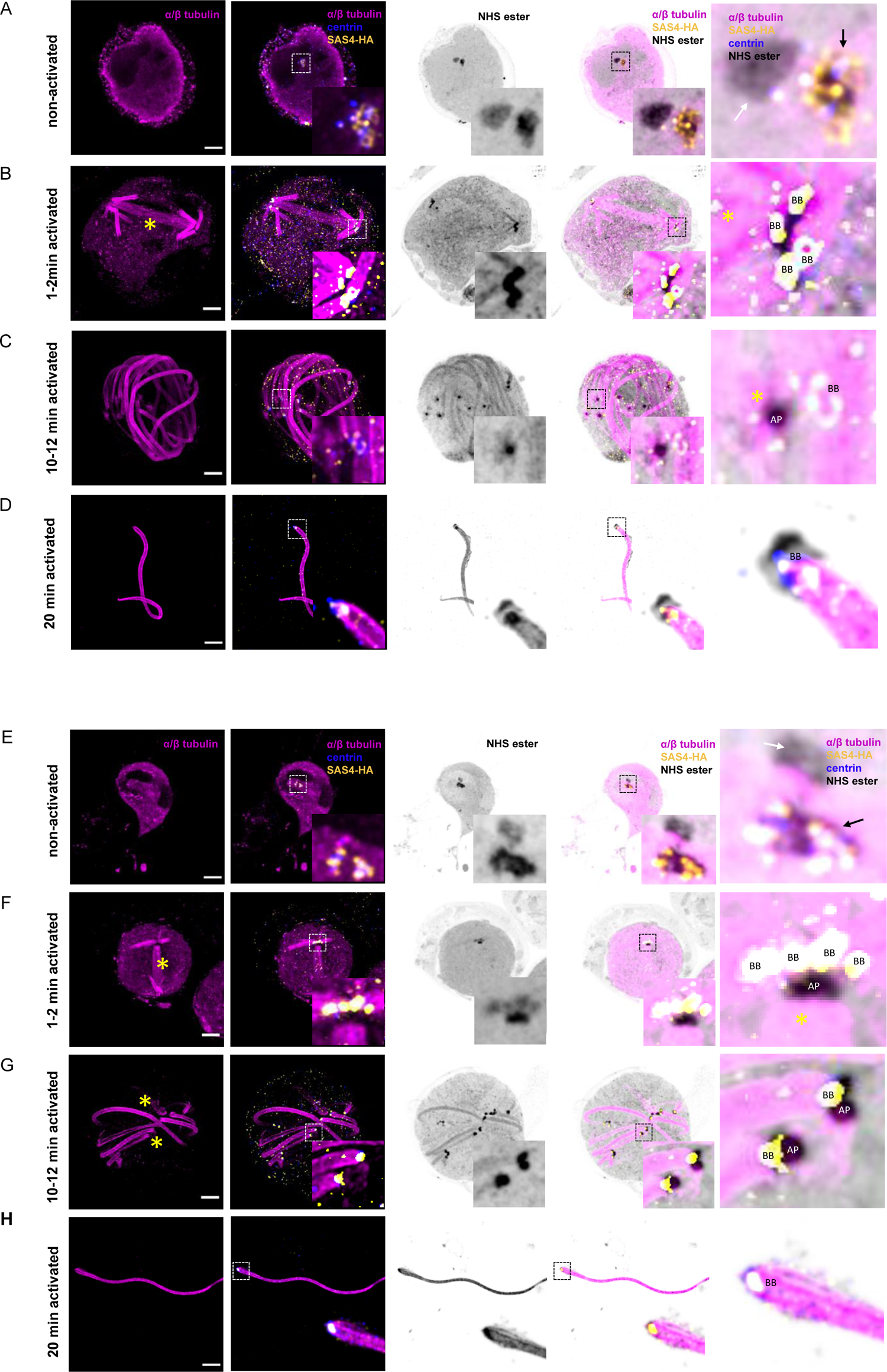
Molecular organisation of the basal body during *P. berghei* microgametogenesis. A-F. Representative full projections of activated *P. berghei* gametocytes. α/β tubulin: magenta; amine reactive groups/NHS-ester: shades of grey; centrin: blue; yellow: SAS4-HA (A-D) or SAS6-HA (E-H). Boxed areas indicate areas shown in insets. Column 5 shows close-up images highlighting the basal bodies (BB), the amorphous MTOC (black arrow), the intranuclear body (white arrow) and acentriolar plaques (AP) stained with NHS-ester and for SAS4-HA (A-D) or SAS6-HA (E-H). Scale bars = 5 µm.

**Figure 8.**
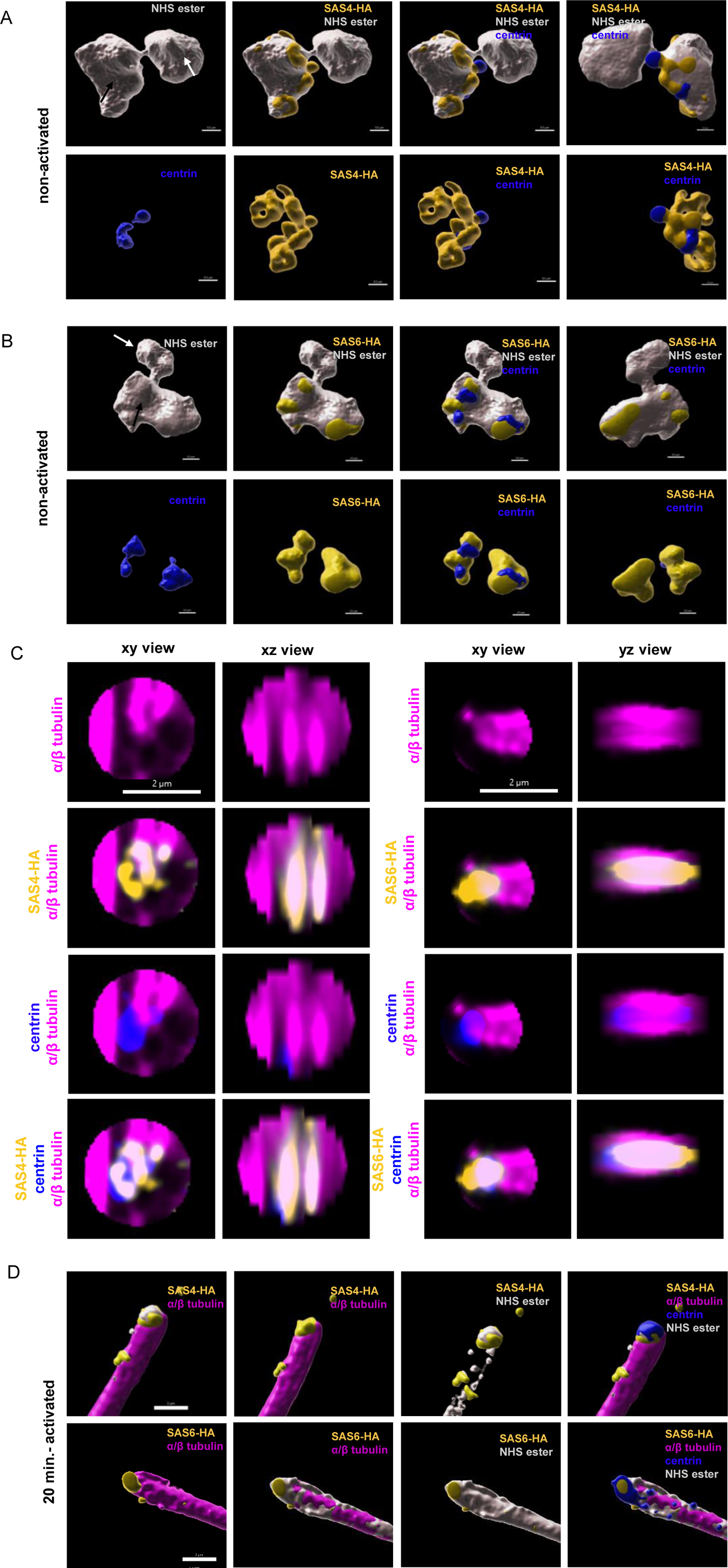
3D reconstruction of the bipartite MTOC during *P. berghei* microgametogenesis. A and B. 3D surface topology of the amorphous MTOC and intranuclear body of non-activated microgametocytes shown in Figure 7. Amine reactive groups/NHS-ester: shades of grey; centrin: blue; yellow: SAS4-HA (A) or SAS6-HA (B); amorphous MTOC: black arrow; intranuclear body: white arrow. Scale bar = 0.5 µm. **C and D.** xy and xz or yz views of individual basal bodies in 10-12 min activated microgametocytes, SAS4-HA (left) and SAS6-HA (right). Scale bar = 2 µm. **E.** 3D surface topology of exflagellated microgametes of figure 7 as shown in A and B. Scale bar = 2 µm.

In 1-2 min activated gametocytes, γ-tubulin was sparsely localised at the extremity of the newly formed basal bodies but was enriched between the NHS-ester dense acentriolar plaque and the nucleation tip of spindle microtubules (Fig. 6B). On the cytoplasmic side, a tetrad of four adjacent basal bodies in front of each acentriolar plaque was visible. SAS4-HA and SAS6-HA were localised to NHS-ester dense basal bodies (Fig. 7B and F). At the proximal end of the axoneme the diameter of the microtubule wall narrowed down from 220 nm to 120 nm following correction with the 4.2x expansion factor. This section may correspond to the disappearance of the two intra-axonemal microtubule singlets [4]. At the base of the axoneme, a 71 nm long centrin cylinder was observed inside the microtubule wall. A slightly longer SAS6-HA cylinder co-localised with centrin. Around the microtubule wall, SAS4-HA displayed a 105 nm long toroid shape at the base of the centrin/SAS6 cylinder (Fig. 8C). In 10-12 min activated gametocytes, a similar organisation was observed with pairs of acentriolar plaques and basal bodies observed at the extremity of fully formed axonemes (Fig. 7C and G). At that stage, a faint γ-tubulin signal could be detected around the basal bodies while a strong signal was observed between the mitotic plate and the mitotic spindles. This suggested a relocalisation from the basal bodies to the acentriolar plaque during gametogenesis (Fig. 6C). In exflagellated microgametes a similar organisation of the basal body was observed (Fig. 7D and H, Fig. 8D)

Of interest, we noted that in the SAS6-HA line, extra SAS6-HA/NHS-ester structures were occasionally observed at the proximal end of single axonemes. A more dramatic phenotype was observed in a SAS6-GFP line, where the NHS-ester dense amorphous MTOC showed an abnormal structure fully labelled with SAS6-GFP. This was associated with aberrant numbers of basal bodies and defective bundling of axonemes (Fig. S1E), as previously observed in a SAS6-KO line [19]. These observations suggested that the accessibility of the SAS6 C-terminus is important for the correct homeostasis of the basal body (Fig. 7G).

### U-ExM reveals early roles of SRPK1 and CDPK4 in the homeostasis of basal bodies and acentriolar plaques

The visualisation of the basal body and the mitotic spindle by U-ExM and bulk proteome labelling opened the possibility to reassess the role of previously well characterised kinases required for the formation of microgametes. We focused on CDPK4 and SRPK1 that were reported to cooperate in the early events of microgametogenesis and respective knock-out lines showed defects in the formation of mitotic spindles [31].

We first focused on the calcium-dependent protein kinase 4 (CDPK4) that is known to play multiple roles during microgametogenesis including entry into S-phase, the formation of mitotic spindles, and assembly of axonemes [32, 33]. We compared expanded CDPK4-KO microgametocytes with their WT counterpart. In non-activated microgametocytes, no obvious difference could be observed (Fig. 9A). Two minutes upon activation by xanthurenic acid, WT microgametocytes displayed the first mitotic spindle with four growing axonemes at each end. In all five analysed CDPK4-KO microgametocytes, both the intranuclear body and the amorphous MTOC remained unchanged as seen by NHS-ester and centrin labelling. As a consequence, no mitotic spindle and no tetrads of axonemes, were observed (Fig. 9B), as previously described [32, 33]. Despite the previously reported defect in DNA replication and the absence of mitotic spindle [32, 33], we observed an increase in the nuclear volume as observed in the WT control. However, 10-12 min after activation, 3-4 bundles of axonemes nucleating from a single MTOC were observed in the 5 mutant microgametocytes. These observations indicated that despite defective reorganisation of the amorphous MTOC, microtubule nucleation was induced but delayed in the absence of CDPK4 (Fig. 9C).

**Figure 9.**
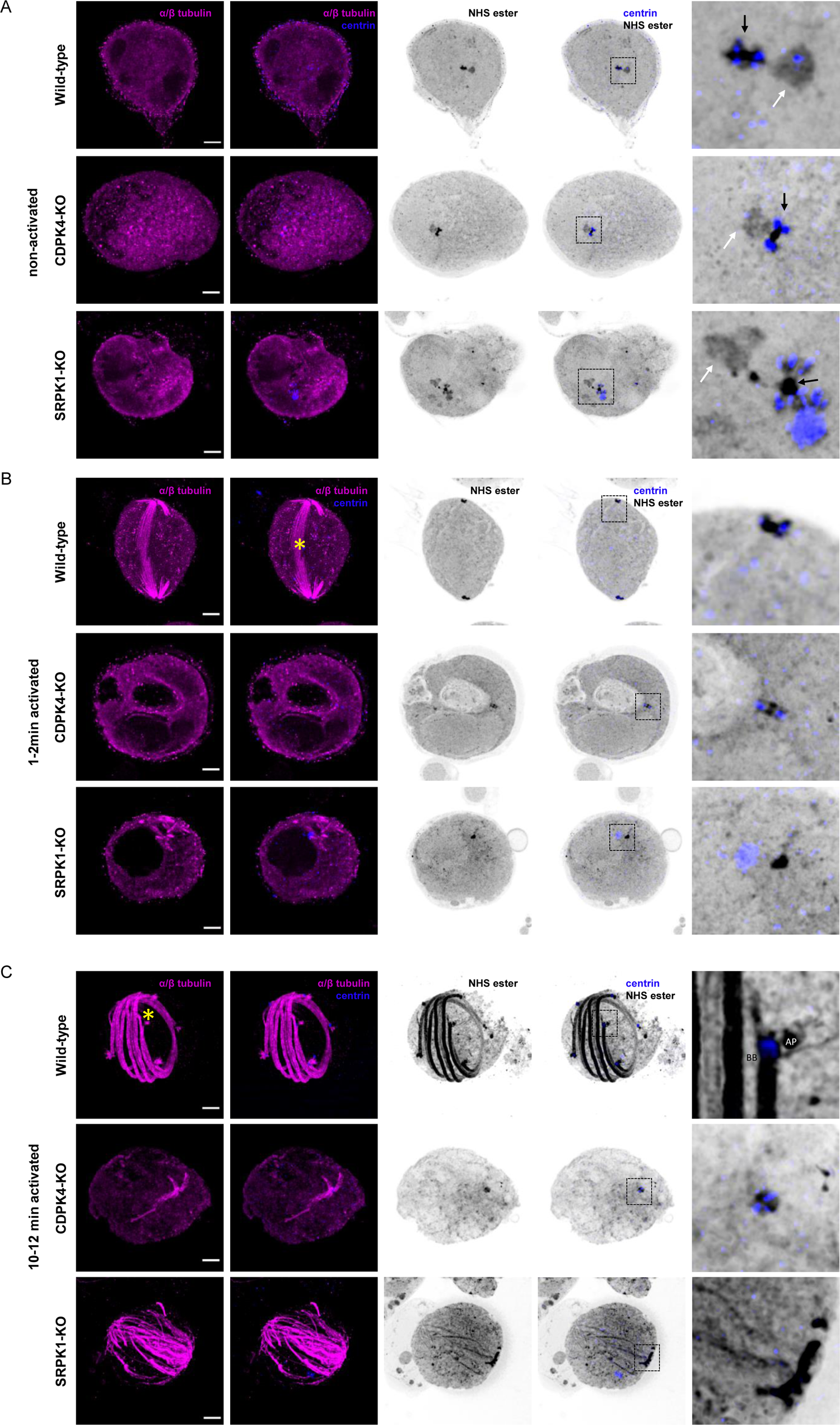
Formation of the amorphous MTOC and its differentiation into functional basal bodies relies on two kinases: CDPK4 and SRPK1. A-C. Representative full projections of (A) non-activated, (B) 1-2 min activated and (C) 10-12 min activated microgametocytes; wild-type (1^st^ row), CDPK4-KO (2^nd^ row) and SRPK1-KO (3^rd^ row). α/β tubulin: magenta; amine reactive groups/NHS-ester: shades of grey; centrin: blue. Boxed areas correspond to close-up images shown in panel 5. Amorphous MTOC: black arrow; intranuclear body: white arrow; basal body: BB; acentriolar plaque: AP; mitotic spindle: yellow asterisk). Scale bars = 5 µm.

In the SRPK1-KO line, more obvious ultrastructural defects were observed in non-activated microgametocytes (Fig. 9A). In non-activated SRPK1-KO microgametocytes, the intranuclear body was absent in 4 out of 5 analysed cells or was detached from the amorphous MTOC in the remaining cell. Additionally, centrin was found in 6-8 NHS-dense spheroids surrounding the amorphous MTOC with one larger centrin positive spheroid in all analysed cells. Upon activation, we did not observe segregation of the basal bodies and the acentriolar plaques (5/5 cells). Despite these early structural defects, however, two tetrads of nucleating axonemes remained in a close position and no mitotic spindle could be observed between these two tetrads (Fig. 9B). Although a faint centrin signal was visible at the proximal end of each basal body, centrin mainly remained in the single large spheroid next to the eight basal bodies (Fig. 9B). Axonemal microtubules did nucleate, but they did not seem to bundle and they remained disorganised in the cytoplasm. In 10-15 min activated microgametocytes, similar defects were still observed, where most basal bodies remained closely clustered. Axonemal microtubules further elongated but lacked a clear organisation with no well-structured axonemes as observed in the parental line (Fig. 9C).

## Discussion

We have recently implemented U-ExM to visualise cytoskeletal structures in various stages of *Plasmodium* parasites [20]. This revealed that a divergent and reduced form of a conoid is present. This organelle is required for host cell invasion by apicomplexan parasites, and was thought to be lost in *Plasmodium*. Here we first combined U-ExM with bulk proteome labelling to investigate further cellular structures in *Plasmodium* gametocytes, the parasite forms that initiate the parasite transmission to the mosquito. NHS-ester staining allowed to resolve numerous structures that were only accessible by EM or super-resolution microscopy such as the sutures between the 13 IMC plates, the osmiophilic bodies, the contour of the nucleus or the host erythrocyte. We additionally observed NHS-ester dense annuli at the extremities of gametocytes that are reminiscent of the apical annuli recently described in *Toxoplasma gondii* tachyzoites [34]. Of particular interest, bulk proteome labelling allowed visualising numerous structures involved in the cell cycle, including the amorphous MTOC and the intranuclear body in non-activated microgametocytes. Upon activation bulk proteome labelling further highlighted the dynamics of these structures as they form cytoplasmic axonemes or mitotic spindles. Pan-ExM further improved the resolution of these structures, however, unlike U-ExM we were unable to label individual proteins, as was done in other cell types [22, 23]. In addition, we found Pan-ExM to be experimentally a lot more demanding than U-ExM. Altogether these observations indicate that the combination of U-ExM with bulk proteome labelling represents an accessible approach to study the ultrastructural context of *Plasmodium* sexual stages with a conventional confocal microscope.

We then exploited the possibility to combine U-ExM with classical immunostaining to clarify the elusive nature of the MTOC in *Plasmodium* gametocytes (Fig. 10). In non-activated gametocytes the NHS-ester dense intranuclear body and amorphous MTOC were connected through the nuclear membrane by proteinaceous filaments. At that stage, the two structures likely host the molecular components required for the formation of the acentriolar plaques and the basal bodies required during microgametogenesis. While none of the proteins investigated could be mapped into the intranuclear body, we show that centrin, γ-tubulin, SAS4-HA and SAS6-HA are present in the amorphous MTOC. At this stage, the localisation of centrin, SAS4-HA and SAS6-HA did not reveal structures resembling basal bodies, as previously suggested by EM [4]. However, the distribution of each protein suggested the presence of defined subdomains within the amorphous MTOC that was surrounded by patches of SAS4. Inside the amorphous MTOC were two to four centrin spheroids surrounded by SAS6 and two elongated γ-tubulin subdomains. This organisation is reminiscent of deuterosomes that serve as platforms to produce basal bodies in a mother centriole-independent manner in multiciliated cells. Deuterosomes are electron dense structures displaying nested subdomains of proteins, including γ-tubulin and SAS6 [35, 36]. Upon activation of microgametocytes, we confirmed the *de novo* assembly of eight basal bodies in the amorphous MTOC through the dynamic localisation of SAS4, SAS6 and centrin. At this stage, in the absence of markers, we were not able to determine whether the intranuclear body rapidly differentiates into eight closely apposed acentriolar plaques upon activation or whether acentriolar plaque division happens sequentially before each round of mitosis. The structural organisation and the composition of the acentriolar plaque thus remains as elusive as its biogenesis. We however show that upon activation, γ-tubulin relocalises rapidly at the interface between the mitotic spindle and the acentriolar plaque, in a pattern that is reminiscent of the multi-layered structures of the yeast spindle bodies [37]. Identification of components of the acentriolar plaques will likely shed light on how the acentriolar plaque is assembled and replicated.

**Figure 10.**
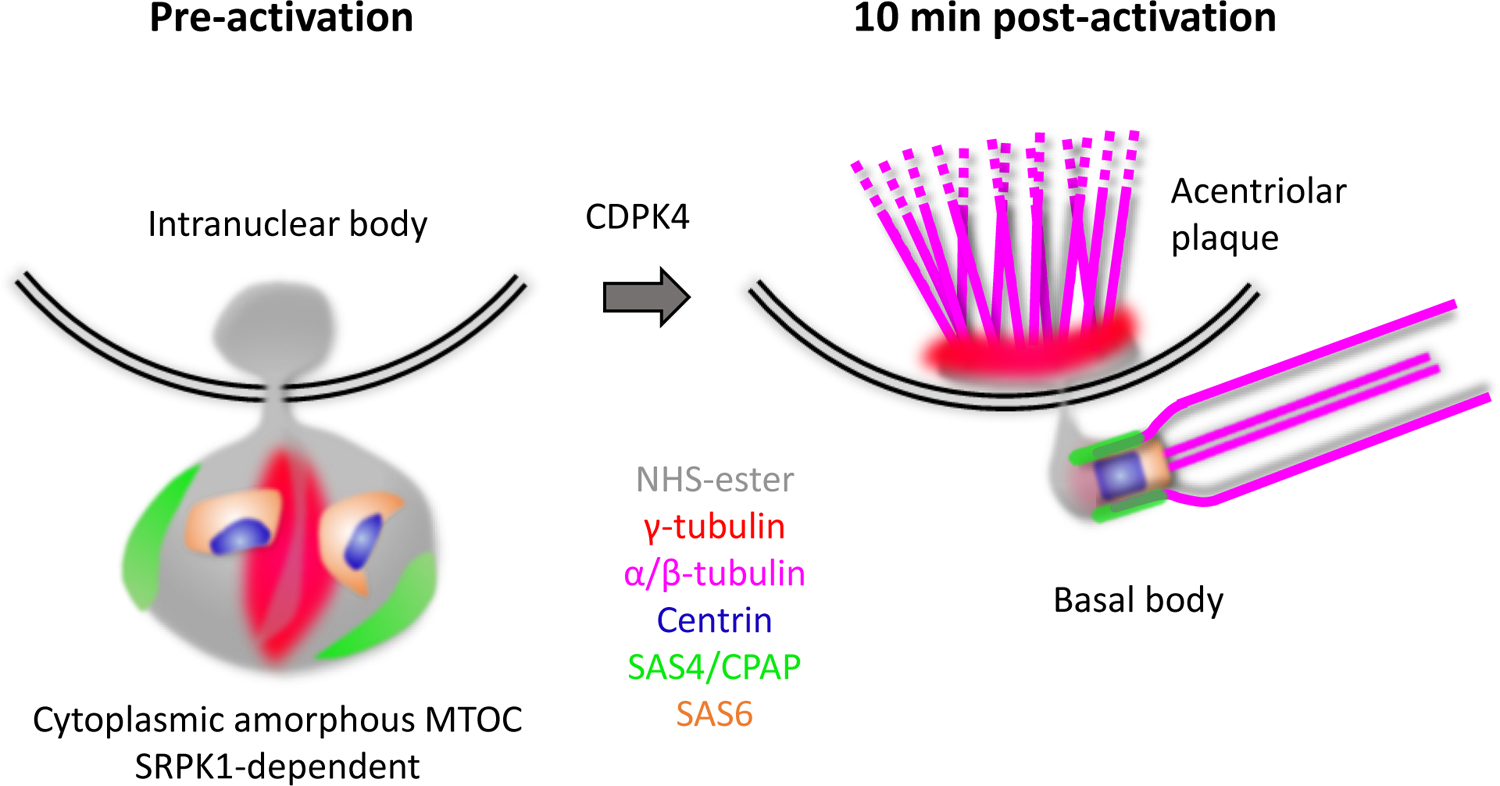
Working model showing the molecular organisation and dynamics of the bipartite *Plasmodium* MTOC coordinating mitosis and axoneme formation during microgametogenesis. In non-activated microgametocyte the intranuclear body is connected to the amorphous MTOC across the nuclear membrane. The amorphous MTOC corresponds to a deuterosome-like structure where components of the basal body and associated proteins localise in distinct or overlapping subdomains. The molecular organisation of the amorphous MTOC and its link with the intranuclear body relies on SRPK1. Upon activation of CDPK4, the intranuclear body gives rise to eight acentriolar plaques that coordinate the assembly of mitotic spindle while the amorphous MTOC differentiates into eight basal bodies to form eight axonemes. γ-tubulin shows dynamic localisation from the amorphous MTOC to the basal bodies and the acentriolar plaques to sequentially nucleate microtubules of the axonemes and mitotic spindles.

Combining U-ExM with localisation of known basal body markers allowed us to refine the structural organisation of this organelle in *Plasmodium*. Upon activation, centrin and SAS6 localised in the proximal lumen of the newly formed microtubule wall, while SAS4 formed a toroid surrounding this structure. In the human centriole, SAS6 is important for the cartwheel formation that lies below Centrin-2. In *Plasmodium*, SAS6 and centrin largely co-localised suggesting that the proximal region and the central core are compacted in a single structure where centrin may not only serve as a scaffolding protein but as a platform for the axoneme nucleation. We also confirm that C-terminal tagging of SAS6 affects the homeostasis of the basal body suggesting the importance of protein-protein interactions in the assembly of this organelle. It remains currently unknown whether the apparently compact structure of the *Plasmodium* basal body is linked to its limited proteome in *Plasmodium* compared with other eukaryotes where up to 100 proteins have been assigned to the centriole [38, 39]. So far, only centrins, SAS6, SAS4 and CEP135 [16–18] have been detected in the *Plasmodium* genome. Interestingly, the latter three proteins are part of the minimal set of proteins required for the nine-fold symmetry of basal bodies [40]. Further functional studies will be required to determine whether additional proteins are involved in the assembly and function of the *Plasmodium* basal body.

The structural organisation of the acentriolar plaque has recently been refined in asexually replicating blood stages [14, 41] that undergo endomitosis but without axoneme nucleation. In these stages, the acentriolar plaque also shows a bipartite organisation with an extranuclear region containing centrin that is linked through a nuclear pore to an intranuclear chromatin-free region harbouring microtubule nucleation sites. In the microgametocyte, a similar organisation of the acentriolar plaque was observed. However, centrin is additionally incorporated into the basal bodies and likely serves as a link to coordinate mitosis with axoneme formation. The shuttling of γ-tubulin from the basal bodies to the acentriolar plaques possibly reflects the sequential nucleation of axonemes and mitotic spindles from this unique bipartite MTOC. In eukaryotes, centrioles are highly conserved and fulfil important cellular functions such as the nucleation of axoneme as a basal body or the organisation of pericentriolar material to form the centrosome. Phylogenetic analyses suggested that the basal body function of the centriole is ancestral whereas its requirement in the organisation of the pericentriolar material emerged later [40, 42]. The coupling of mitosis with axoneme formation in *Plasmodium* gametocytes may reflect a transitory state in the evolution of a basal body function. It also shows functional similarities with the basal body-dependent segregation of the mitochondrial or kinetoplast DNA during the cell cycle of *Trypanosoma brucei* [43]. In the absence of axoneme requirement in the *Plasmodium* asexual blood stages, it is possible that the expression of basal body components was lost in these stages but that the acentriolar plaque retained the bipartite structural scaffold to fulfil mitosis.

Kinase activity has long been known to be crucial for the regulation of basal body homeostasis. A central kinase for basal body biogenesis is the polo-like kinase 4 (PLK4) in humans or related orthologous kinases in other eukaryotes [44]. PLKs were proposed to be present in the cenancestor of eukaryotes but lost in some lineages [42] including *Plasmodium* [45, 46]. It was suggested that in these lineages, either the basal body duplication does not require kinase activity or that other kinases fulfil that role. Here we analysed with U-ExM two previously characterised kinase mutants, SRPK1 and CDPK4, in which mitotic defects during microgametogenesis were observed [31, 33, 45]. A SRPK1-KO line produced morphologically normal microgametocytes despite substantial deregulation of phospho-dependent pathways prior to activation [20]. Here U-ExM reveals structural defects of the amorphous MTOC in terminally differentiated SRPK1-KO gametocytes, confirming the role of SRPK1 prior to activation. Deletion of *srpk1* is linked with misincorporation of centrin in the amorphous MTOC leading to defects in basal body segregation and axoneme bundling. While U-ExM did not allow us to define precisely subdomains in the amorphous MTOC of WT parasites, the mislocalisation of centrin in the SRPK1-KO line further supports the presence of a defined molecular architecture within the amorphous MTOC. Deletion of *cdpk4* was previously shown to lead to a strong reduction in the formation of mitotic spindle and axonemes [15, 32]. We further show that both defects are linked to an early block in the *de novo* formation of the eight basal bodies. Despite this marked defect, the formation of short axonemal-like structures is observed at later time-points suggesting that other regulatory mechanisms are at play to induce microtubule formation upon activation of microgametocytes. Interestingly, CDPK4 and SRPK1 are not essential in replicative asexual blood stages [45]. We here show they are associated with integrity of the amorphous MTOC that is not expressed during these stages. It is thus tempting to speculate that kinases important for the homeostasis of the acentriolar plaque will play important roles in both asexual and sexual blood stages.

In this work, the high resolution of U-ExM has allowed us to gain new insights into the structural and molecular organisation of an atypical bipartite MTOC coordinating mitosis and axoneme formation in a non-model eukaryote. Deeper characterisation of its molecular composition and its homeostasis may in the future help highlighting general principles underlying the evolution and organisation of MTOC in eukaryotes.

## Methods

### Key resource table

| Key resource table |  |  |  |  |
| --- | --- | --- | --- | --- |
| Reagent type or resource | Designation | Source or reference | Identifiers | Additional information |
| Cell line<br>*P. falciparum* | iGP2 | Boltryk et al, 2020 |  |  |
| Cell line<br>*P. berghei* | ANKA 2.34 | Billker et al, 2004 |  |  |
| Cell line<br>*P. berghei* | GEX1-HA | Ning et al, 2013 |  |  |
| Cell line<br>*P. berghei* | SAS4-HA | This study |  |  |
| Cell line<br>*P. berghei* | SAS6-HA | This study |  |  |
| Cell line<br>*P. berghei* | CDPK4KO | Billker et al, 2004 |  |  |
| Cell line<br>*P. berghei* | SRPK1-KO | Tewari et al, 2010 |  |  |
| Antibody | Centrin mouse (20H5) | Merck Millipore | 04-1624 | U-ExM: 1:500 |
| Antibody | α-tubulin Guinea pig | Unige antibody platform | AA345 | U-ExM: 1:250 |
| Antibody | β-tubulin Guinea pig | Unige antibody platform | AA344 | U-ExM: 1:250 |
| Antibody | γ-tubulin Rabbit | Sigma | T5192 | U-ExM: 1:500 |
| Antibody | HA Rat (3F10) | Roche | 11815016001 | U-ExM 1:250 |
| Antibody | Actin Mouse | Herm-Gotz et al, 2002 |  | U-ExM 1:3 |
| Antibody | anti-mouse Alexa 488 | Invitrogen | A11001 | U-ExM: 1:400 |
| Antibody | anti-guinea pig 488 | Invitrogen | A11073 | U-ExM: 1:400 |
| Antibody | anti-rat 488 | Invitrogen | A11006 | U-ExM: 1:400 |
| Antibody | anti-rabbit Alexa 405 | Invitrogen | A31556 | U-ExM: 1:400 |
| Antibody | anti-mouse Alexa 405 | Invitrogen | A31553 | U-ExM: 1:400 |
| Antibody | anti-guinea pig Alexa 647, | Invitrogen | A21450 | U-ExM: 1:400 |
| Chemical | Atto 594 NHS-ester | Merck | 08741 | 10 µg/mL |
| Chemical | Hoechst 33342 | Invitrogen | H3570 | 5:1000 |

### Ethics statement

All animal experiments were conducted with the authorisation number GE/58/19, according to the guidelines and regulations issued by the Swiss Federal Veterinary Office.

### *P*. *berghei* maintenance and gametocyte production

*P*. *berghei* ANKA strain-derived clone 2.34 [32] together with derived transgenic lines GEX1-HA [47], SAS4-HA, SAS6-HA, CDPK4-KO [32] and SRPK1-KO [45] were grown and maintained in CD1 outbred mice as previously described [48]. Six- to ten-week-old mice were obtained from Charles River Laboratories (France), and females were used for all experiments. Mice were specific pathogen free (including *Mycoplasma pulmonis*) and subjected to regular pathogen monitoring by sentinel screening. They were housed in individually ventilated cages furnished with a cardboard mouse house and Nestlet, maintained at 21 ± 2°C under a 12-hour light/dark cycle, and given commercially prepared autoclaved dry rodent diet and water ad libitum. The parasitemia of infected animals was determined by microscopy of methanol-fixed and Giemsa-stained thin blood smears.

Gametocyte production and purification was performed as previously described [48]. Parasites were grown in mice that had been phenyl hydrazine-treated 3 days before infection. One day after infection, sulfadiazine (20 mg/L) was added in the drinking water to eliminate asexually replicating parasites. For gametocyte purification, parasites were harvested in suspended animation medium (SA; RPMI 1640 containing 25 mM HEPES, 5% fetal calf serum [FCS], 4 mM sodium bicarbonate, pH 7.20) and separated from uninfected erythrocytes on a Histodenz cushion made from 48% of a Histodenz stock (27.6% [w/v] Histodenz [Sigma/Alere Technologies, Germany] in 5.0 mM TrisHCl, 3.0 mM KCl, 0.3 mM EDTA, pH 7.20) and 52% SA, final pH 7.2. Gametocytes were harvested from the interface. Gametocytes were activated in RPMI 1640 containing 25 mM HEPES, 4 mM sodium bicarbonate, 5% FCS, 100 μM xanthurenic acid, pH 7.4).

### Generation of targeting constructs and parasite transfection

In order to generate the transgenic SAS4-HA and SAS6-HA lines, library clones PbG02_A-10a01 and PbG03-39b05 from the PlasmoGEM repository (http://plasmogem.sanger.ac.uk/) were used to generate HA tagging vectors, respectively.

Sequential recombineering and gateway steps were performed as previously described [49, 50] using RecUp and RecDown primers. Insertion of the GW cassette following gateway reaction was confirmed using primer pairs GW1 x QCR1 and GW2 x QCR2. Oligonucleotides used in this study are listed in Table S1. The modified library inserts were then released from the plasmid backbone using NotI. The HA targeting vectors were transfected into the 2.34 parasite line. Schizonts for transfection were obtained and parasite transfection was performed as previously described [48].

### P. falciparum culture

*In vitro* cultures to produce *P. falciparum* gametocytes using the iGP2 line was performed as described in [24]. Briefly, the parasite line was grown in human erythrocytes in RPMI-1640 medium with glutamine (Gibco), 0.2% sodium bicarbonate, 25 mM HEPES, 0.2% glucose, 5% human serum, and 0.1% Albumax II (Life Technologies). The medium was complemented with 2.5 mM D-Glucosamine (D-GlcN) to restrict the overexpression of the sexual commitment factor GDV1 under the control of a *glmS* riboswitch. At ring stage parasites, D-GLcN was removed to trigger the expression of GDV1 to initiate gametocytogenesis. Gametocyte maturation was allowed for 10 to 15 days. For the first 6 days following induction, 50 mM GlcNAc was added to the culture medium to eliminate asexual parasites. Parasite development was monitored daily by Giemsa staining.

### U-ExM

Synchronized stage specific *P. falciparum* iGP line cultures were pelleted and fixed in 4% formaldehyde for 20 minutes. Sample preparation of *P. berghei* parasites for U-ExM was performed as previously described [20], except that 4% formaldehyde (FA) was used as fixative. Fixed samples were then attached on a 12 mm round Poly-D-Lysine (A3890401, Gibco) coated coverslips for 10 minutes. Thereafter the following steps were performed: 1. To add anchors to proteins, coverslips were incubated for 5 hours in 1.4% formaldehyde (FA)/ 2% acrylamide (AA) at 37°C, 2. Gelation was performed in ammonium persulfate (APS)/Temed (10% each)/Monomer solution (23% Sodium Acrylate; 10% AA; 0,1% BIS-AA in PBS) for 1 hour at 37°C. 3. Denaturation was performed for 1 hour and 30 minutes at 95°C. 4. Gel expansion and antibody labelling. After denaturation, gels were incubated in ddH2O at room temperature for 30 minutes. Next, gels were incubated in ddH2O overnight for complete expansion. The following day, gels were washed in PBS twice for 15 minutes to remove excess of water. Gels were then incubated with primary antibodies at 37°C for 3 hours, and washed 3 times for 10 minutes in PBS-Tween 0.1%. Incubation with the secondary antibody was performed for 3 hours at 37°C followed by 3 washes of 10 minutes in PBS-Tween 0.1% (all antibody incubation steps were performed with 120-160 rpm shaking at 37°C). Directly after antibody staining, gels were incubated in 1 ml of 594 NHS-ester (Merck: 08741) diluted at 10 μg/mL in PBS for 1 hour and 30 minutes at room temperature on a shaker. The gels where then washed 3 times for 15 minutes with PBS-Tween 0.1% and expanded overnight in ultrapure water. Overnight, a second round of expansion was done in water before imaging. 5. Sample mounting and imaging: 1cm x 1cm gel pieces were cut from the expanded gel and attached on 24 mm round Poly-D-Lysine (A3890401, Gibco) coated coverslips to prevent gel from sliding and to avoid drifting while imaging. The coverslip was then mounted on a metallic O-ring 35mm imaging chamber (Okolab, RA-35-18 2000-06) and imaged.

### Pan-ExM

The pan-ExM procedure was performed as previously published [23]. Briefly, samples were fixed in 4% formaldehyde and attached on a 12 mm round Poly-D-Lysine (A3890401, Gibco) coated coverslips for 10 minutes. Thereafter the following steps were performed. 1. Addition of anchors to proteins: coverslips were incubated for 5 hours in anchoring solution, 1.4% FA/ 2% AA mix at 37°C 2. 1^st^ Gelation: gelation was performed in a gelation chamber, using APS/Temed/ (19% Sodium Acrylate; 10% AA; 0.1% N,N-(1,2-dihydroxyethylene) bis-acrylamide-DHEBA-in PBS) for 1 hour at 37°C. 3. Denaturation: the gel was carefully removed from the gelation chamber and placed in the denaturation buffer for 1 hour and 30 minutes at 76°C. 4. First expansion: after denaturation, a piece of the 1 cm x 1 cm gel was cut and placed in ultra-pure water for about 2 hours; water was changed 2 times for the first 30 min. This leads to an expansion factor ranging from 4x to 4.5x. 5. Re-embedding in neutral hydrogel/2^nd^ gel: 2 cm x 2 cm gels were cut and used for further downstream processes as bigger gels are difficult to handle and require a bigger volume of reagents. Water was replaced by the second gelling solution (APS/Temed/10% AA, 0.05% DHEBA) and incubated for 3 x 20 min at room temperature on a shaker. Thereafter the gel was placed in a humid gelation chamber and incubated at 37°C for 2 hours. 5. Addition of anchors to proteins: the gel was removed from the gelation chamber and placed in anchoring solution (1.4% FA/ 2% AA) at 37°C for 5 hours. 6. 3^rd^ gelation: The gel was washed three times in PBS and incubated three times with the third gelation solution APS/Temed/ (19% Sodium Acrylate; 10% AA; BIS-AA in PBS) on ice on a shaker. After the third incubation, the gel was placed on a glass slide and covered on top with a coverslip (22 mm x 22 mm), placed in a humidified chamber and incubated at 37°C for 2 hours. 7. Dissolution of 1^st^ and 2^nd^ gel: the gel was removed from the humidified chamber and incubated with 0.2 M NaOH solution in a 6-well plate for 1 hour at room temperature on a shaker. After 1 hour, the solution was removed, rinsed with PBS and transferred to a beaker with PBS and washed 3 to 4 times for 30 min each or until the pH changed to 7.4. 8. NHS-ester staining: the gel was placed in a 6 well plate and incubated in 594 NHS-ester (Merck, 08741) diluted at 10 μg/mL in 3 ml PBS for 1 hour and 30 minutes at room temperature on a rocking platform. The gels were then washed three times for 15 minutes with PBS-Tween 0.1%. 9. 2^nd^ expansion: gels were place in a beaker filled with ultra-pure water and incubated for 4-6 hours; water was changed every 30 minutes for the first hour of incubation. 10. Sample mounting and imaging: this step was performed exactly as explained above for U-ExM.

### Image acquisition, analysis and processing

Images were acquired on Leica TCS SP8 microscope with HC PL Apo 100x/ 1.40 oil immersion objective in lightning mode to generate deconvolved images. System optimized Z stacks were captured between frames using HyD as detector. Images were processed with ImageJ, LAS X and Imaris 9.6 software. Imaris software was used for 3D surface reconstruction and xy, xz and yz representations.

## Supplementary material

**Figure S1. Generation and characterisation of SAS4-HA, SAS6-HA and SAS6-GFP transgenic lines. A-D.** Genetic modification scheme and genotyping of the transgenic lines. **E.** SAS6-GFP microgametocytes display an abnormally shaped amorphous MTOC densely stained for SAS6-GFP. This does not seem to affect mitosis but axoneme formation and arrangement is compromised. α/β tubulin: magenta; amine reactive groups/NHS-ester: shades of grey; centrin: blue; SAS6-GFP: yellow. Scale bars = 5 µm.

**Movie S1.** 3D surface topology reconstruction of the microgametocyte shown in Figure 3B. α/β tubulin: magenta; amine reactive groups/NHS-ester: grey.

**Table S1.**
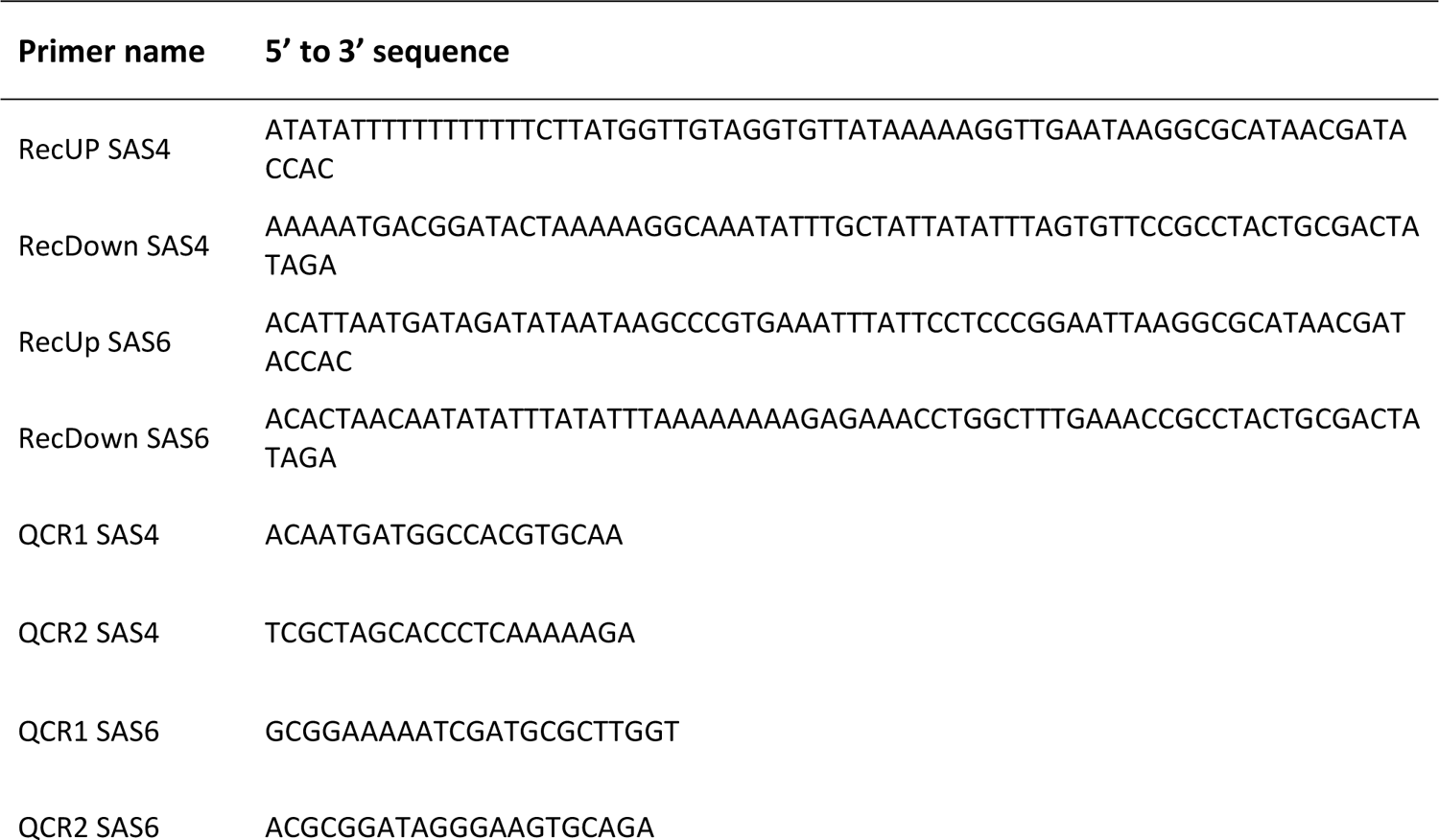
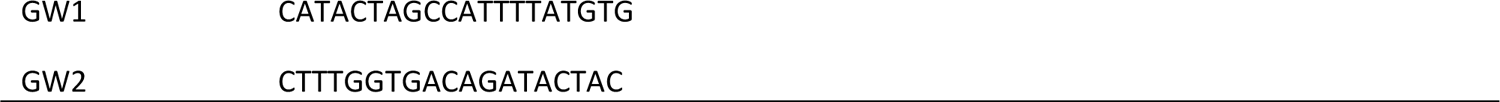
Oligonucleotides used in this study.

## Author Contributions

MB conceived and supervised the project. MB and RR designed the experiments and wrote the manuscript. RR performed the experiments and analysed the data.

## Acknowledgments

We would like to thank Till Voss (Swiss Tropical and Health institute) for generously sharing the *P. falciparum* iGP2 line. We thank Virginie Hamel and Paul Guichard for insightful discussions on the biology of the centriole, we also thank Eloïse Bertiaux and Vincent Louvel from their laboratory for discussions on the U-ExM and Pan-ExM protocols. We thank Dominique Soldati-Favre for sharing reagents and general discussions on the biology of apicomplexan parasites, as well as Oscar Vadas for critical reading of the manuscript. We also thank Natacha Klages for technical assistance. We finally thank the excellent service at the bioimaging core facility at the Faculty of Medicine of the University of Geneva. This work was supported by the Swiss National Science Foundation (SNSF) 31003A_179321 to MB. MB is an INSERM and EMBO young investigator.

## Data and materials availability

All data needed to evaluate the conclusions in the paper are present in the paper and/or the supplementary materials. Additional data or reagents are available from authors upon request. Correspondence and requests for materials should be addressed to M.B.

