## Supplementary figures and images for "Ultrastructure expansion microscopy of *Plasmodium* gametocytes reveals the molecular architecture of a microtubule organisation centre coordinating mitosis with axoneme assembly"

### Figure S1

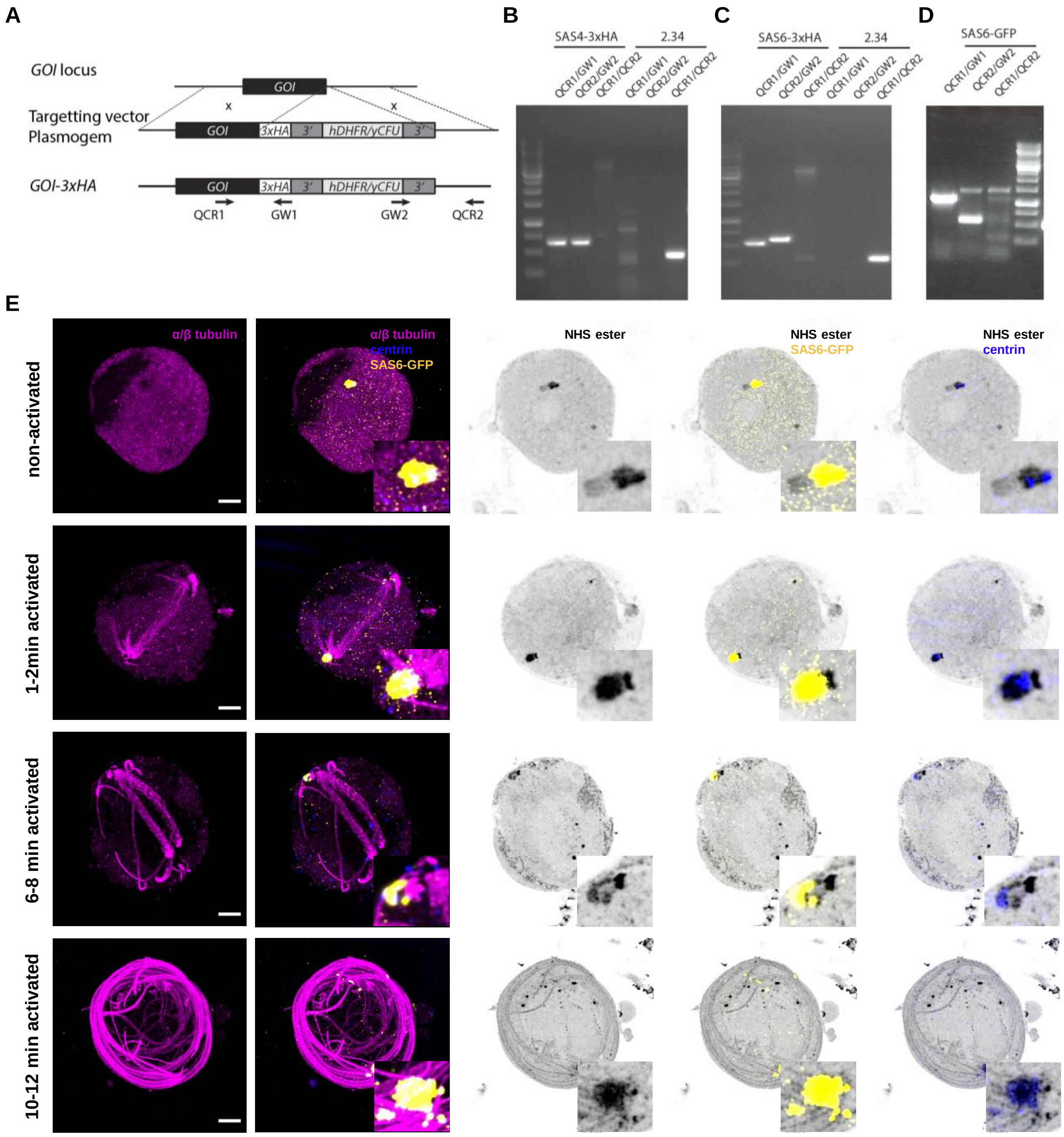
